# Synaptic input sequence discrimination on behavioral time-scales mediated by reaction-diffusion chemistry in dendrites

**DOI:** 10.1101/106567

**Authors:** Upinder Singh Bhalla

**Author notes:** Lead Contact: Upinder S. Bhalla.

## Abstract

Sequences of events are ubiquitous in sensory, motor, and cognitive function. Key computational operations, including pattern recognition, event prediction, and plasticity, involve neural discrimination of spatio-temporal sequences. Here we show that synaptically-driven reaction-diffusion pathways on dendrites can perform sequence discrimination on behaviorally relevant time-scales. We used abstract signaling models to show that this selectivity arises when inputs at successive locations are aligned with, and amplified by, propagating chemical waves triggered by previous inputs. We incorporated biological detail using sequential synaptic input onto spines in morphologically, electrically, and chemically detailed pyramidal neuronal models. Again, sequences were recognized, and local channel modulation on the length-scale of sequence input could elicit changes in neuronal firing. We predict that dendritic sequence-recognition zones occupy 5 to 20 microns and recognize time-intervals of 0.2 to 5s. We suggest that this mechanism provides highly parallel and selective neural computation in a functionally important time range.

## Introduction

Activity sequences have long been recognized as a fundamental constituent of neural processing. Lorente de No suggested that reverberatory activity sequences in small networks could sustain activity (Lorente de No, 1938). Hebb’s idea of cell assemblies suggested that ensembles of cells encoded a particular neuronal concept, but also that there was sequential activation within the group of cells forming the assembly (Hebb, 1949).

Many sensory systems process sequential stimuli, and these are typically mapped to ensembles of sequentially active neurons (Bouchard and Brainard, 2016; Broome et al., 2006; Carrillo-Reid et al., 2015). Deeper in the brain, hippocampal place cells represent a higher-order cognitive map of space, yet here too sequences occur when the animal moves through spatial locations represented by the place cells (Wilson and McNaughton, 1994). The hippocampus exhibits other forms of sequential activity in the form of fast replay events (Jadhav et al., 2012; Wilson and McNaughton, 1994) and stimulus-bridging activity that emerges during associative learning (MacDonald et al., 2011) and trace conditioning (Modi et al., 2014). Thus there are numerous neural correlates both of sequences, and of processing steps that recognize them.

The idea of synfire chains examines conditions for self-sustaining sequential activity to occur in multiple layers of a network (Abeles, 1982). This is non-trivial, as excessive activation can lead to runaway epileptiform activity, whereas insufficient activation causes decay of the activity wave (Kumar et al., 2008; Mehring et al., 2003). Neural networks that can recognize such sequences are well-established. For example, events may be run through a delay line, so that the oldest event is delayed more, the next event less, and so on, so that they all arrive at the neural network at the same time. Thus the time-sequence is flattened in time and classical attractor networks can recognize patterns in the sequence (Tank and Hopfield, 1987). Time-varying attractor networks have also been shown to be able to implement and recognize sequences (Lee, 2002). Time-invariance is a desirable feature of sequence recognition circuits, since the same order of events may take place at different speeds. Recursive networks using supervised learning, and short-term synaptic plasticity have been shown to be implement time-invariance (Barak and Tsodyks, 2006; Goudar and Buonomano, 2015; Laje and Buonomano, 2013).

While networks can carry out sequence recognition, they make limited use of the rich dynamics of biological neurons. The theory of Heirarchical Temporal Memory builds on the idea of sequence-recognizing neurons and networks as a way to perform complex temporal computation (George and Hawkins, 2009; Hawkins and Ahmad, 2016). Even the ability to recognize simultaneous closely-localized synaptic input has interesting computational implications (Hawkins and Ahmad, 2016), The current study extends this to patterns of input both in time and space.

One of the first biophysical proposals for subcellular sequence recognition was made by Rall, who showed theoretically that synaptic events propagate down the dendritic tree with a small delay. By timing synaptic inputs to coincide with this delay, Rall predicted that ordered input should yield a larger response than reversed input (Rall, 1964). A stronger version of this mechanism was experimentally demonstrated by Branco et.al. (Branco et al., 2010) who used glutamate uncaging to provide sequential input along a pyramidal neuron dendrite. They showed that NMDA receptor amplification of ‘inward’ sequences (distal to proximal) gave rise to about 40% larger somatic depolarization than ‘outward’ sequences, for rapid (~40ms) sequences. However, many interesting neural sequences occur at slower time-scales.

At least three attributes should converge for single neurons to achieve and report sequence recognition in the noisy context of neural activity. First, the neurons should recognize inputs coming in the correct order in space and time. Second, the selectivity for the correct input over scrambled input and background noise should be strong enough for there to be reliable discrimination. Third, recognized sequences should trigger activity changes either by way of changed firing, or by way of plasticity. In the current study we use theory and reaction-diffusion modeling to show how the first condition may be achieved, and develop detailed multi-scale models to address the second and third.

## Results

As the setting for this study we considered a network in which ensembles of neurons are active in sequence, for example, place cells in the hippocampus as an animal moves along a linear track (Figure 1). We assumed that a single axon from each of these ensembles projects onto spines located on a short stretch of dendrite, in the same spatial and temporal order as the activation of the ensembles. We first explored the discrimination of such sequential input using abstract chemistry in a one-dimensional diffusion geometry. We then mapped the discrimination mechanism to a signaling pathway modeled as mass-action reaction-diffusion chemistry in a similar cylindrical geometry, but ornamented with dendritic spines. We then tested discrimination when we embedded mass-action chemistry into a morphologically and electrically detailed neuronal model. Finally we asked if ion channel modulation by sequence discrimination chemistry in small dendritic zones, could affect neuronal firing.

**Figure 1.**
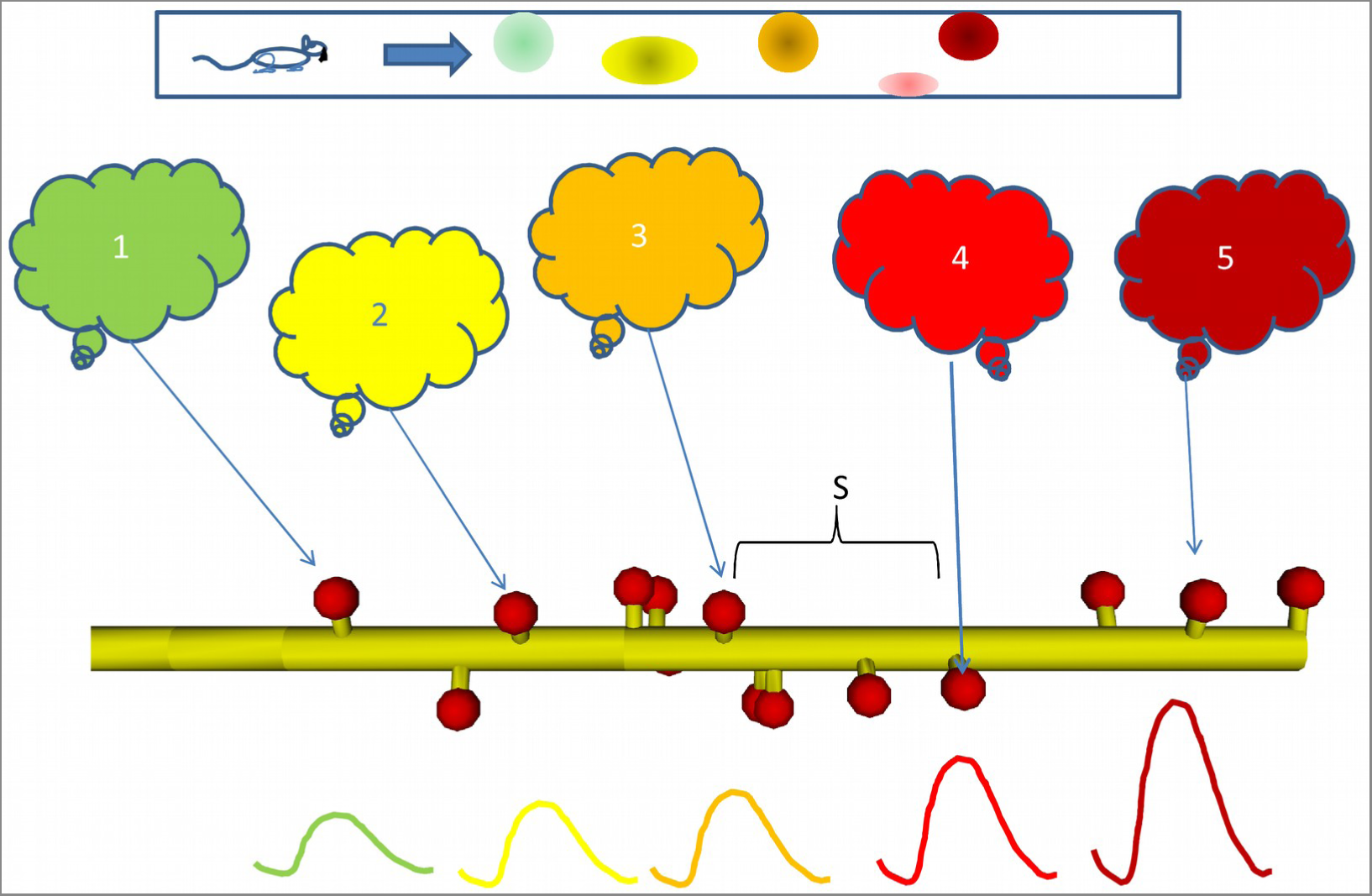
Sequential activity, from behavior to dendrite. Top: Linear arena in which a rat moves, with indicated locations of 5 place cells (color coded). The place cells are representatives of 5 neuronal ensembles (colored clouds), each active in one of the 5 locations on the arena. The neuronal ensembles each send a single axonal projection in spatial order to a small dendritic segment, with an average spacing *S* between connected spines. Note that spines need not be immediately adjacent to each other. Below, buildup of reactant following input activity with the appropriate timing, corresponding to the rat moving at a speed which the dendritic chemistry recognizes.

### Abstract reaction-diffusion systems support sequence recognition

We first analyzed the requirements for chemical reaction-diffusion systems to achieve selectivity for spatially and temporally ordered inputs. In doing these calculations we utilized ‘spherical cow’ models of chemistry: highly reduced formulations with just two diffusive state variables A and B, interpreted as molecules undergoing 1-dimensional diffusion (Figure 2A). The reaction system received input from a third molecule, designated as Ca (Figure 2B).

**Figure 2.**
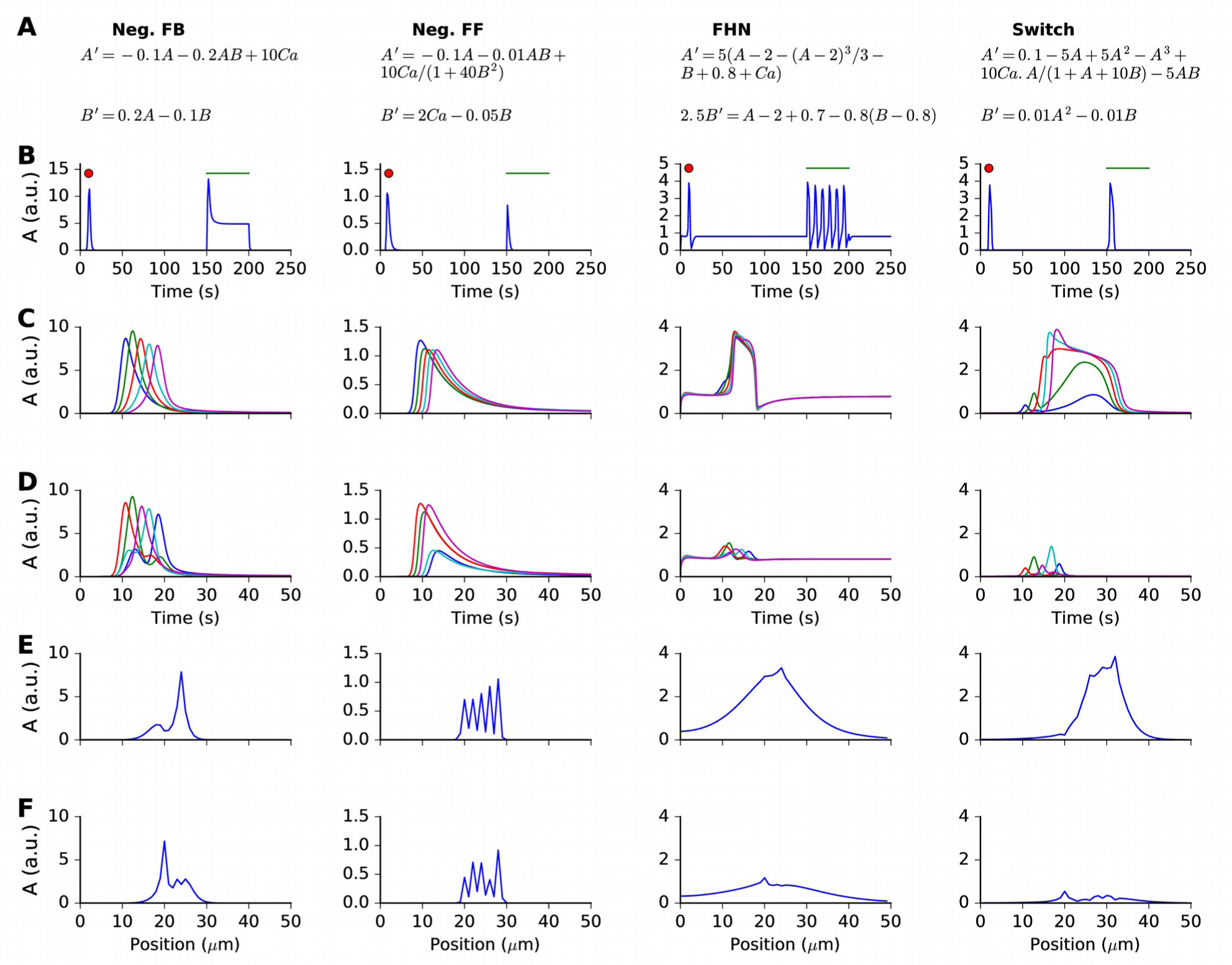
Responses and sequence selectivity of four abstract models involving molecules A and B. Columns are the respective models. A. Model equations. B. Response of point (non-spatial) model to a 1s wide Gaussian input of Ca^2+^ (red dot) and a steady pulse of Ca ^2+^ (green line). Input amplitudes: 1, 10, 0.4, and 1 respectively. C-F. Response of molecule A in one-dimensional reaction-diffusion form of model to sequential input at 5 locations. C. Time-course of response at 5 locations to ordered input. D. Time-course of response at 5 locations to scrambled input. Note that the FHN and Switch models have much lower responses to scrambled as compared to ordered input. The Neg. FF model has a lower response at two of its locations. E, F: Snapshot of spatial profile of response to ordered (E) and scrambled (F) input. Snapshot is at time 18.4, 14.2, 16.3 and 18.4 s respectively.

Our criterion for sequence recognition was that ordered inputs should elicit high total activity of a signaling molecule A, whereas scrambled inputs should elicit low levels of activity. Total activity here means the sum of activity of A over the duration and spatial extent of the stimulus.

We examined four abstract reaction systems: nonlinear feedback inhibition; nonlinear feedforward inhibition; an explicit propagating wave solution based on the FitzHugh-Nagumo equation (Fitzhugh, 1961; Nagumo et al., 1962) and a fast positive, slow negative feedback system exhibiting state switching followed by turnoff (Figure 2A). In all cases molecule B inhibited A.

To investigate the temporal responses of these ‘reaction’ systems, we first delivered impulse (red dot) and step-function (green line) input to non-spatial versions of the models (Figure 2B). Each model exhibited a large impulse response. Two of the models (feedforward inhibition and state-switching) exhibited only a transient response to the sustained input. The feedback model, as expected, had a transient strong response followed by a shallow sustained response. The FitzHugh-Nagumo model, again as expected, oscillated.

We then implemented 1-dimensional reaction-diffusion versions of these models (Figure 2). We delivered Ca^2+^ stimuli at 5 equally spaced points on the reaction system. Each Ca^2+^ stimulus followed a Gaussian time-profile:

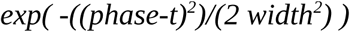

Here *phase* defines the timing of each input, *t* is time, and *width* is the duration of the Ca^2+^ pulse. The only difference between the sequential and scrambled stimuli was their order.

The results of these simulations are compared in Figure 2C–F. First, we found that the simple feedback inhibition model showed no input sequence-dependent difference in total activity. This can be seen in terms of time-response at the 5 stimulus points (Figure 2C,D), and also in terms of spatial profile of A at the end of the stimulus (Figure 2E,F).

Then, we observed that the feedforward inhibition model showed a small amount of selectivity (Figure 2C–F). This arose because the A response was diminished at two of the input points when the input was scrambled.

The FitzHugh-Nagumo (FHN) model was strongly selective, with a large buildup of response only when the input was sequential (Figure 2 third column). We interpret this as arising when the propagating wave from the FHN equation arrived at successive input points just when the input was also present.

The switching model was the most selective (Figure 2 last column). This was because it built up to a large, sustained response lasting several seconds, provided the successive inputs were in the same position as the diffusively propagating activity of A.

Thus this set of simulations showed that several reaction-diffusion like systems were capable of sequence selectivity. Strong selectivity emerged from a supralinear buildup of responses when diffusively propagating activity was aligned with successive inputs, and suppression due to inhbition when the alignment was off.

### Reaction systems select for distinct speeds and length-scales of sequential input

We devised a scalar metric of the degree of sequential ordering, and used this metric to see how the different reaction systems were tuned to different spatial and temporal intervals. We plotted the ordinal position of the input against the ordinal value of its arrival time. Thus a perfect sequence would arrive at positions [0,1,2,3,4] at times [0,1,2,3,4]. We used the quantity

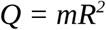

as the measure of how ordered the sequence was. Here, *m* = slope and *R* = regression coefficient of best-fit line. With this measure, the sequence [0,1,2,3,4] has Q = 1; [4,3,2,1,0] has Q = −1, and [4,0,2,1,3] has Q = −0.001.

We then compared our reaction readout A with the metric Q, to see how well each of our four reaction systems could discriminate sequence order (Figure 3 A–D, first column). The results were as expected from the individual runs from Figure 2. Specifically, the inhibitory feedback system had poor tuning, there was a small amount of selectivity for the feedforward inhibition case, and extremely strong selectivity for the FitzHugh-Nagumo and switching cases. Note that the modeled reaction-diffusion system was spatially symmetric and did not distinguish between forward and backward sequences. An analysis of symmetry breaking arising from incorporation of biological detail is out of the scope of the current paper.

**Figure 3.**
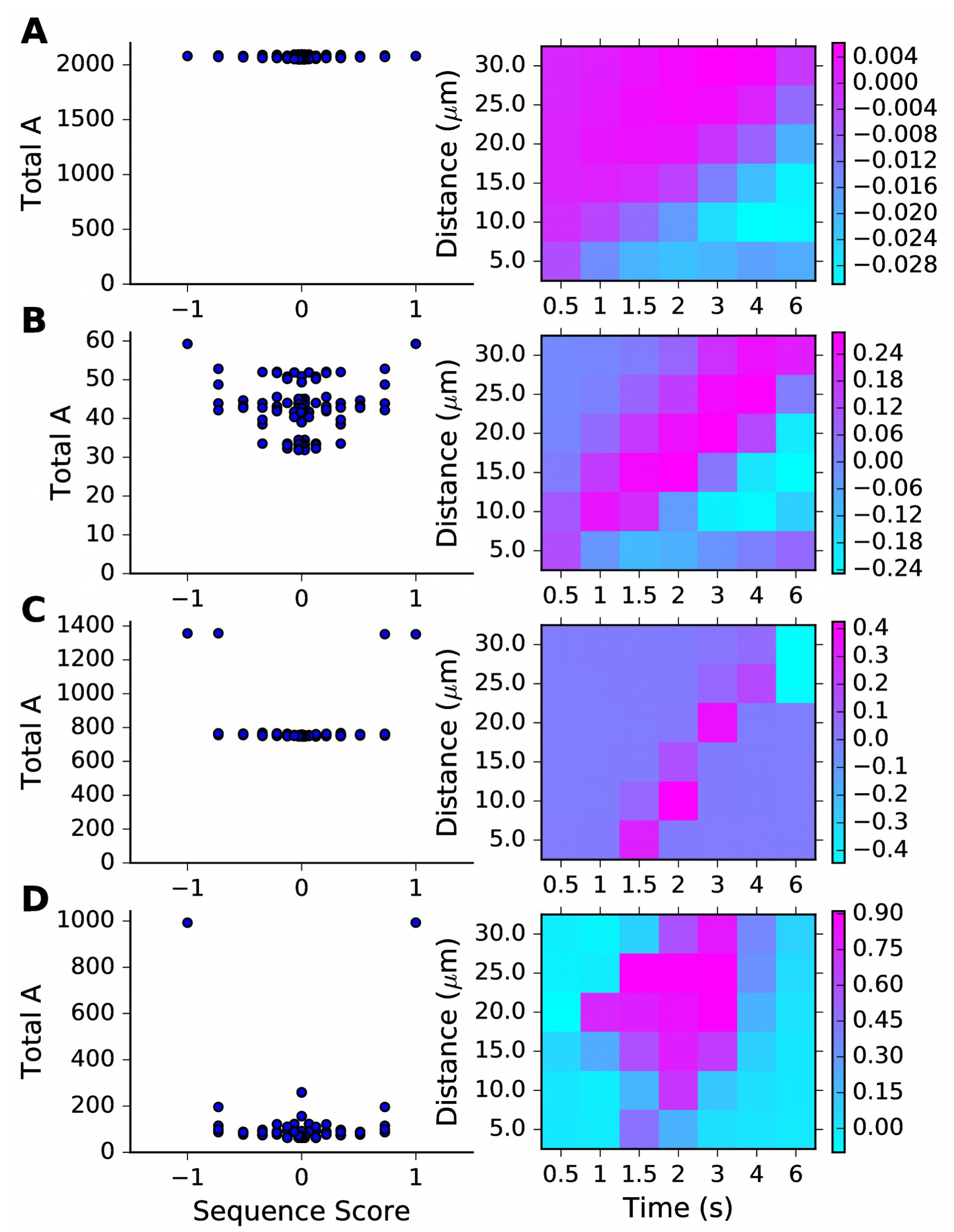
Sequence selectivity dependence on space and time intervals. Left column: scatter plots of chemical system selectivity (measured as total activation of molecule A over time and space) against linearity of input sequence. Right column: Matrix of selectivity score Q as a function of total length of stimulus zone and interval time between successive stimuli. A: Feedback model. B: Inhibitory feedforward model. C: FitzHugh Nagumo model. D: Switching model.

We next asked how selective each reaction system became, when the inputs were delivered at different time and space intervals. We devised a metric for reaction network sequence selectivity, based on a comparison of responses to sequential vs scrambled inputs:

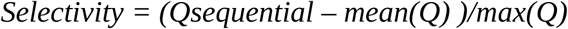

Thus, to compute *Selectivity* at each value of input timing and spacing, we carried out simulations of the model for all possible non-repeating patterns of input to obtain a comprehensive set of *Q* values. We repeated these calculations for each point in the matrix of timing and spacing values (Figure 3 second column). As expected, the feedback inhibition model had low selectivity. The feedforward inhibition model showed a diagonal band of weak selectivity such that rapid stimuli in close proximity as well as slower stimuli at greater distances were preferred. The FitzHugh-Nagumo case was much more strongly selective, but also had a similar diagonal band. In both these cases we interpret this as there being an intrinsic propagation speed of the chemical activity wave, and if the inputs were timed and spaced accordingly, the response built up. The switching model was different, in having a large diffuse zone of strong selectivity centred around (2 seconds, 4 microns) between stimulus points.

Thus, each of the three sequence-selective reaction systems had a preferred range of stimulus timings and spacings.

### Sequence speed selectivity scales with reaction rates and diffusion constants

As a further analysis on our abstract models, we asked how sequence speed selectivity scaled with rates and diffusion parameters. This scaling is important because it determines the spatial extent of the sequence selective zones that we propose to exist in the dendritic tree. It is also important as it defines the time-scales of sequential events that may be recognized by such reaction-diffusion mechanisms.

We first tested the most straightforward assumption, that sequence propagation speed was proportional to chemical rate and diffusion constants. To do this we scaled all rates and the diffusion constants by the same factor, and reduced the stimulus width by the same factor. We ran the same grid of stimulus space and time intervals as above. As expected, we found that as the rates were increased, the best tuning was at shorter time intervals but roughly the same spatial intervals. (Figure 4 A–E). However, different stimulus strengths were needed to obtain strong tuning in these runs. We therefore also examined dependence of tuning on stimulus strength, and observed that the tuning zone also depended on stimulus strength (Figure 4 F, G). This is because the selectivity occurs when the stimulus is strong enough that the ordered sequence leads to pathway activation, but no so strong that the stimulus overrides the inhibitory reactions and produces indiscriminate activation.

**Figure 4.**
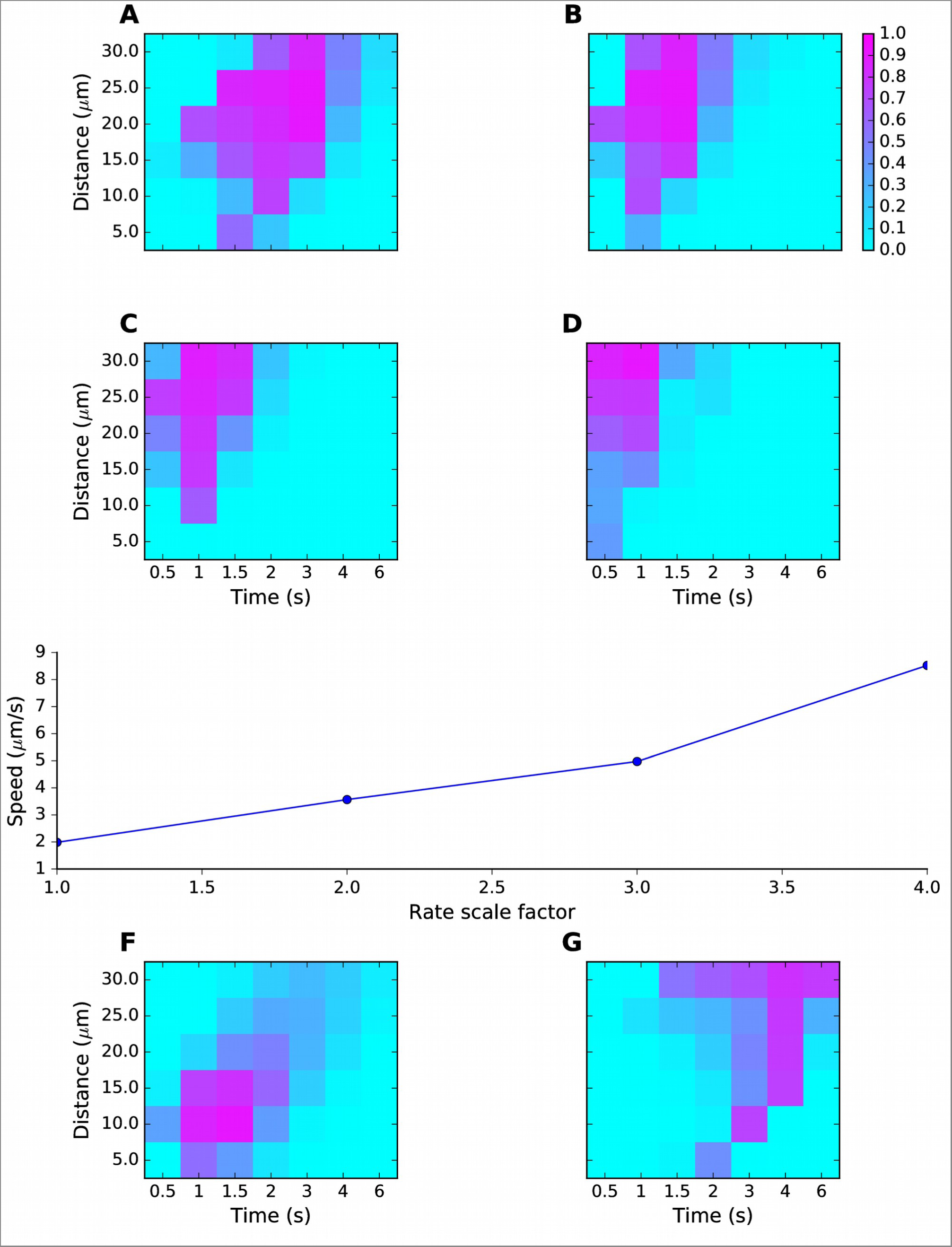
Sequence tuning ranges in space and time. A-D: tuning for rates of 1, 2, 3 and 4 times basal, respectively. E: Plot of speed of sequence as a function of the rate scale factor. F, G: Preferred space/time of tuning shifts with stimulus amplitude. F: 90% basal. G: 110% basal.

Thus, by varying rates for reactions and diffusion, we were able to achieve good sequence selectivity over a broad range of time-intervals (0.5-4 s between stimuli) and spatial intervals (2-6 mm between stimuli).

### Mass-action reaction-diffusion system based on MAPK feedback discriminates sequences

We next asked if above design principles for sequence selectivity could be applied to a mass-action reaction-diffusion system. We based our mass-action model on a published, reduced model of MAPK feedback and bistability (Bhalla, 2011). The model already exhibited the switch-like turnon behavior of our abstract ‘switch’ model. The other key aspect of the abstract model was delayed turnoff of activity. There are several known inhibitory feedback turnoff mechanisms for the MAPK pathway (Lake et al., 2016). We therefore added a MAPK-activated protein phosphatase to cause delayed turnoff of the kinase. The other changes we made to the published MAPK model were a) to remove the synaptic signaling leading to receptor turnover in the spine and b) to add diffusible CaM as a Ca^2+^ buffer. With the exception of active PKC, Raf, and the ion channels, all molecules were diffusible. The model schematic is shown in Figure 5A, and the full model specification is presented in supplementary material. This model formulation is a semi-quantitative map to the far more detailed and tightly constrained models of this pathway in the literature (Bhalla and Iyengar, 1999; Resat et al., 2003). Ca^2+^ stimuli were delivered in the PSD and diffused into the dendrite (Methods). Model responses to brief (1s) and square step function (100s) Ca^2+^ inputs are shown in Figure 5B,C, both measured in the dendrite. As expected, there is a strong switch-like turnon to Ca^2+^ stimuli, followed by delayed inhibition. We ran this model in a cylindrical geometry ornamented with spines at ~1.1 micron intervals (methods, Figure 5D). We provided sequential and scrambled Ca^2+^ input to 5 of the spines, and observed good sequence selectivity (Figure 5E,F).

**Figure 5.**
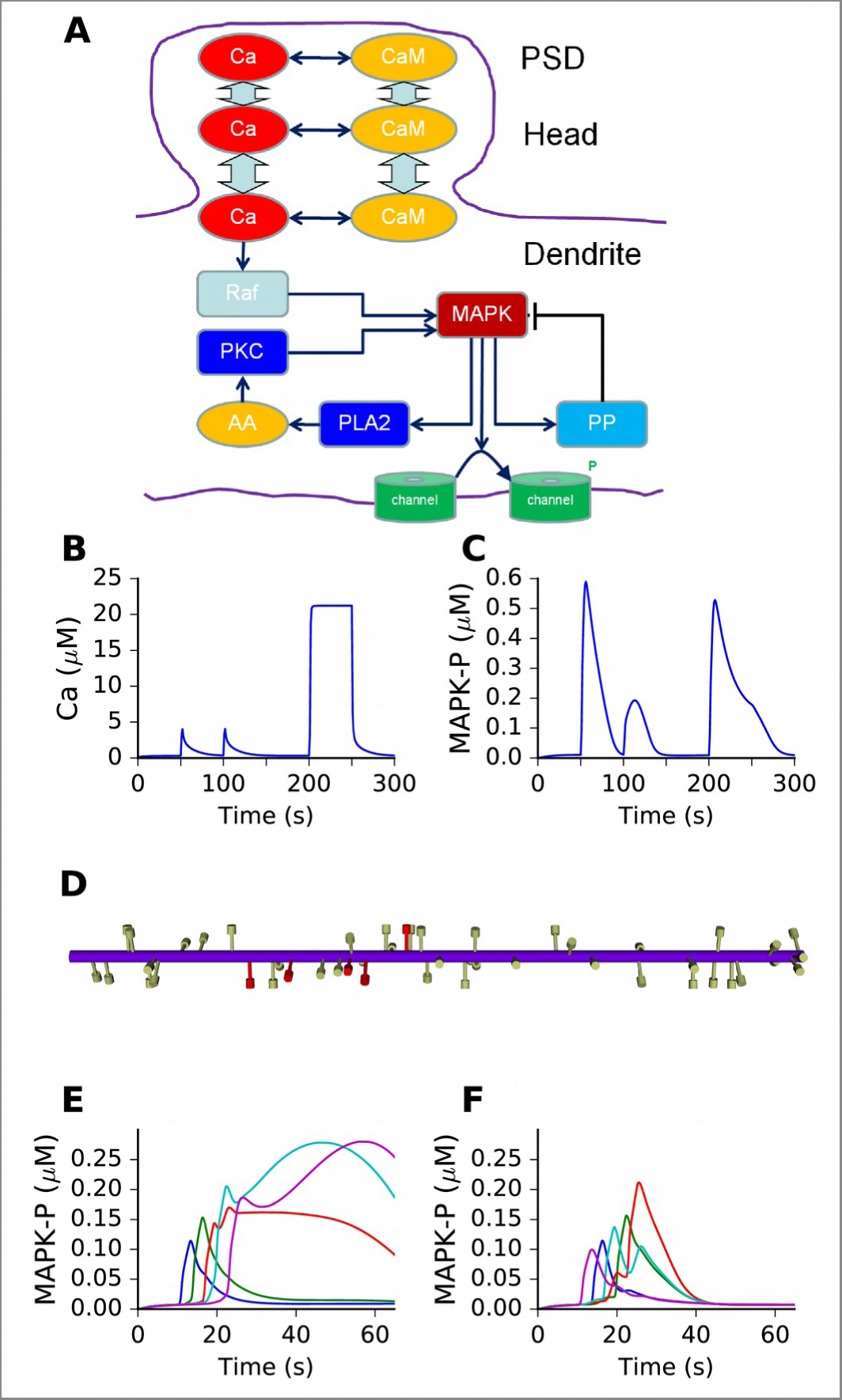
Mass action model of a MAPK switch exhibits sequence selectivity. A. Schematic of model chemistry. Black arrows indicate binding or activation reactions, plunger indicates inhibition, and bent arrow indicates enzyme activity. Broad cyan arrows indicate diffusion of Ca^2+^ and CaM between PSD, spine head, and dendrite. CaM buffers the incoming Ca^2+^ from the PSD. B. Ca^2+^ responses seen at the dendrite, to input delivered at PSD. PSD stimulus was two pulses of 1 s, 160 μM Ca^2+^, separated by 50 s. After a delay of 100 s, a step of 160 μM Ca^2+^ was delivered for 50 s. C. MAPK-P response to the Ca^2+^ stimulus. Note strong inhibition of the response to the second pulse, and rapid decline of response to the 50 step stimulus. D. Geometry of 1-D reaction-diffusion model, with 49 spines. Dendrite diameter was 1 μm and length was 60 μm. The 5 stimulated spines are in red. E. Response of system to sequential input (01234) on spines spaced ~ 3 microns apart, at intervals of 3 seconds, starting on the 11^th^ spine from the left. Each input was delivered to the PSD of the stimulated spine, at 160 μM, for 2.9 s. MAPK-P was recorded under each of the stimulated spines F. Response to scrambled input, in the order 40312. The response to scrambled input had a smaller amplitude and lasted for a shorter time.

Thus we showed that a mass-action reaction-diffusion system exhibited sequence selectivity when it had switch-like turnon with delayed turnoff by negative feedback. This configuration had been predicted by the abstract models.

### Sequence selectivity works in a biologically detailed multiscale neuronal model with noisy input

We then asked whether biochemical sequence recognition would ‘work’ in the more complex context of active neurons receiving background activity in a network. We brought in biological detail at the following levels: a) Model neuron morphology based on anatomical reconstructions (>3000 segments), including >5700 spines spaced at ~1 micron. b) Voltage-gated ion channels distributed throught the cell, based on published models. c) Background glutamatergic Poisson synaptic input at 0.1 Hz on all spines. d) Background GABAergic synaptic input with an 8 Hz theta modulation on proximal dendritic compartments. Due to the background synaptic input, the model exhibited theta-modulated subthreshold oscillations, with spiking activity at ~1 Hz (Figure 6B). e) Stimuli in the form of spike trains arriving on AMPA and NMDA receptors on subsets of spines. f) Ca^2+^ dynamics following synaptic ion flux, including diffusion between spine and dendrite, and buffering by calmodulin, in all spines and dendrites. g) The reduced MAPK model described above, was distributed throughout the dendritic tree in ~6000 diffusive compartments (Figure 6 A). h) Chemical calculations in the spines were carried out using a stochastic method (Gillespie Stochastic Systems Algorithm, methods) for the runs used to calculate selectivity.

**Figure 6.**
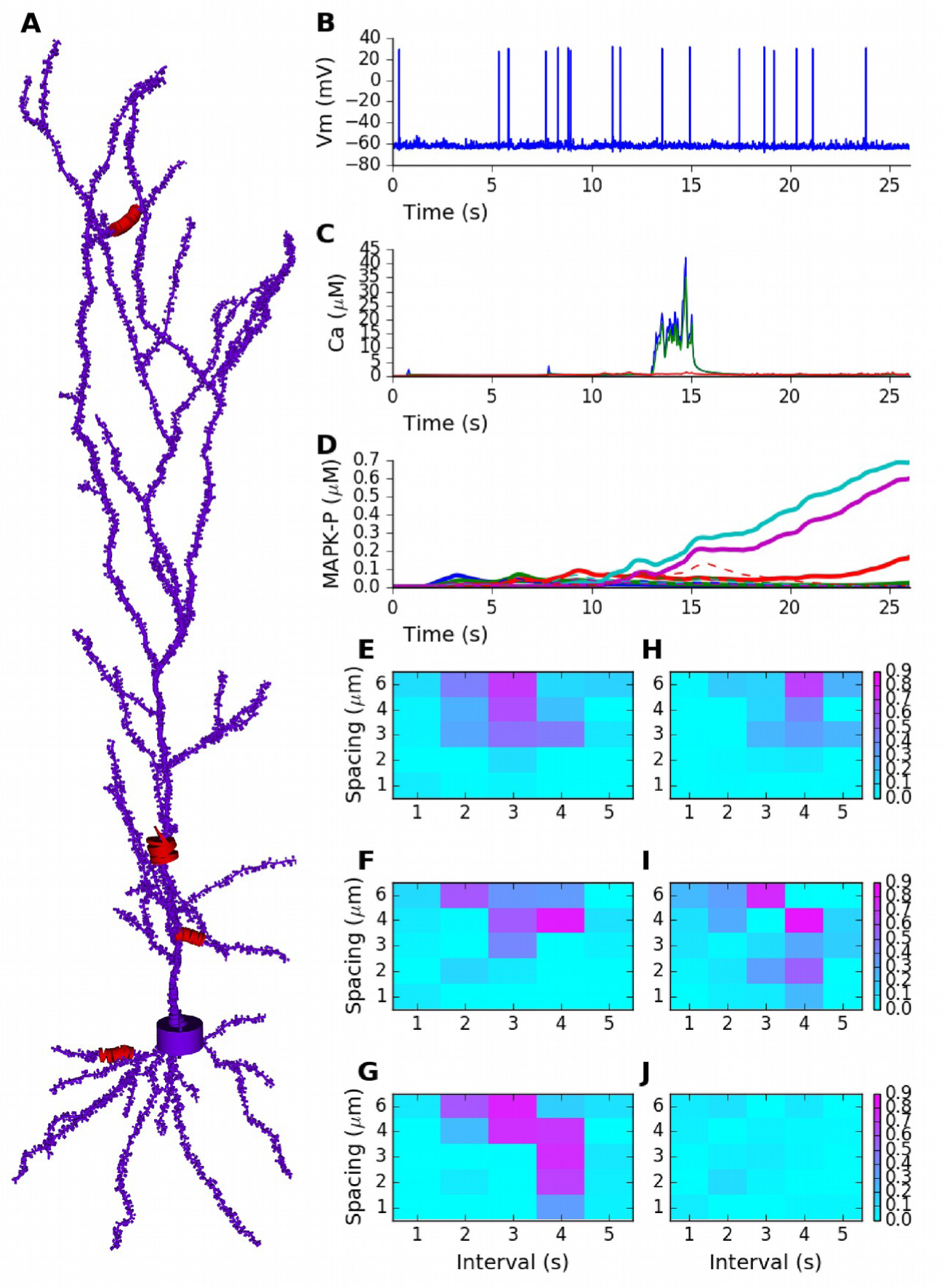
Sequence selectivity in detailed electrical+chemical signaling model. A. Morphology of model. Dendrite diameters are scaled up by 2, and spine diameters and lengths by 4 for visualization. Stimulus was given in 4 zones on the cell, indicated in red and by diameters scaled up by 10x. B. Example somatic intracellular potential and spike train of model neuron. C. Ca^2+^ responses to Poisson synaptic volley at mean rate 20 Hz, measured in PSD (blue), spine head (green) and dendrite (red). D. Sequence selectivity in the distal apical zone. Heavy solid lines are P-MAPK levels under the stimulated spines, for sequential input. Strong buildup occurs for three of the spines. Dashed lines are corresponding P-MAPK levels for scrambled input. Only the sequential input responses lead to build up. E-G: Matrix of sequence selectivity in basal dendrite zone, for different input spacing in time and space. Three different runs are shown, with the same morphology but different random number seeds for the background and stimulus synaptic input. H: Selectivity in proximal oblique dendrite zone, using 40% larger spine dimensions. I: Selectivity in distal apical dendrite using 20% larger spines. J: Selectivity matrix in proximal primary apical dendrite shows no sequence selectivity even with 40% larger spines.

We first ran a direct comparison of a sequential input train [01234] compared with a scrambled train with sequence [40312]. We picked time and space intervals of (3s, 4 microns) based on preliminary calculations for tuning. We delivered the input sequence on a basal dendritic branch (Figure 6A). We selected a set of 5 spines, spaced at ~3 microns. Each spine was stimulated with a Poisson spike burst lasting 2 s at 20Hz on the AMPA and NMDA receptors. With these parameters, the [Ca^2+^] reached ~40 mM in the PSD, ~35 mM in the spine head, and ~1 mM in the region of dendrite immediately below the stimulated spine (e.g., Figure 6C). Similar Ca^2+^ levels were obtained when stimuli were delivered in other regions of the dendrite. The exception was the primary apical dendrite, in which dendritic Ca^2+^ only reached ~0.5 μM. This is an expected outcome of diluting out the Ca^2+^ arriving from the spine, since the diameter of the primary dendrite was ~2 μM as compared to ~1.0 μM in the other stimulus regions. With these parameters, we observed strong selectivity in MAPK activity for sequential stimuli as compared to scrambled in the basal dendrite (Figure 6 D, solid lines for sequential vs. dashed for scrambled).

We then asked how this multiscale model responded to a range of time and space intervals in each of the stimulated zones. Because the calculations were expensive, we used a modified version of the selectivity metric (equation 3) in which we computed outcomes of only 12 stimulus patterns rather than the exhaustive 120 possible permutations (methods). For efficiency, we delivered the stimulus patterns simultaneously in the 4 zones of the cell (Figure 6A). Similar tuning was observed in test simulations where stimuli were delivered only in a single zone.

We observed strong sequence selectivity in the basal dendrite zone, for a restricted range of space and time intervals, as expected from the earlier calculations (Figure 6 E–G). We repeated the calculations using different random seeds in order to generate different background and stimulus spiking input to the model neuron. We found that the responses of individual runs, and hence selectivity, were somewhat noisy (Figure 6 E–G).

We next investigated whether the response selectivity was manifested in different regions of the cell. We found that there were indeed strongly sequence selective responses in oblique dendrites and distal apical dendritic regions, but these required stronger stimuli (Figure 6 H, I). In these runs we had to increase the diameters of spine heads and spine shafts by 40% and 20% respectively to obtain sufficent dendritic Ca^2+^ influx, and hence selectivity. The reference spine models had a shaft of 1 μm length × 0.2 μm diameter, and a head of length and diameter 0.5 μm. We did not observe sequence recognition in the primary apical dendrite (Figure 6J). Even the strongest stimuli applied in this location elicited only small Ca^2+^ elevations, with weak downstream MAPK responses, and no pattern selectivity.

Thus we showed that intracellular signaling pathways exhibited selectivity for ordered synaptic input, even when considerable morphological and electrical detail and noisy input were incorporated. Selectivity became noisier with this additional detail, and different zones of the cell required different synaptic efficacies to achieve selectivity. With the current parameters the selectivity was on the time-scale of seconds, with successive inputs spaced apart by a few microns.

### Local dendritic channel modulations may influence cellular firing

There are multiple possible outcomes of local sequential activity-triggered chemical signaling. These include synaptic plasticity, dendrite remodeling, and electrical changes through channel modulation. For the purposes of the current study we focused on the third, namely changes in electrical activity, since this defines how the neuron may report the presence of a sequence to other cells in the neuronal network. Similar brief, strong synaptic input on small sections of distal dendrites has been reported to cause strong changes in activity of striatal spiny neurons by triggering a transition to an ‘up’ state (Plotkin et al., 2011).

We utilized the same morphologically detailed pyramidal neuron model as above, and asked if local channel modulations over the length scale of about 20 microns could alter somatic spiking. We applied these channel modulations to the same four regions of the cell which we had tested for sequence discrimination. We examined modulation of KA, Na, leak, NMDAR and AMPAR. We estimated the statistics of firing following these modulatory changes through repeated simulations of the control condition (100 repeats) and each modulation condition (40 repeats), each with different random number seeds. We found that modulation of the Na channel, and increase in leak conductance in the apical and primary dendrites could increase firing rates by large factors (Figure 7A, B). In initial runs receptor modulation had no effect.

**Figure 7.**
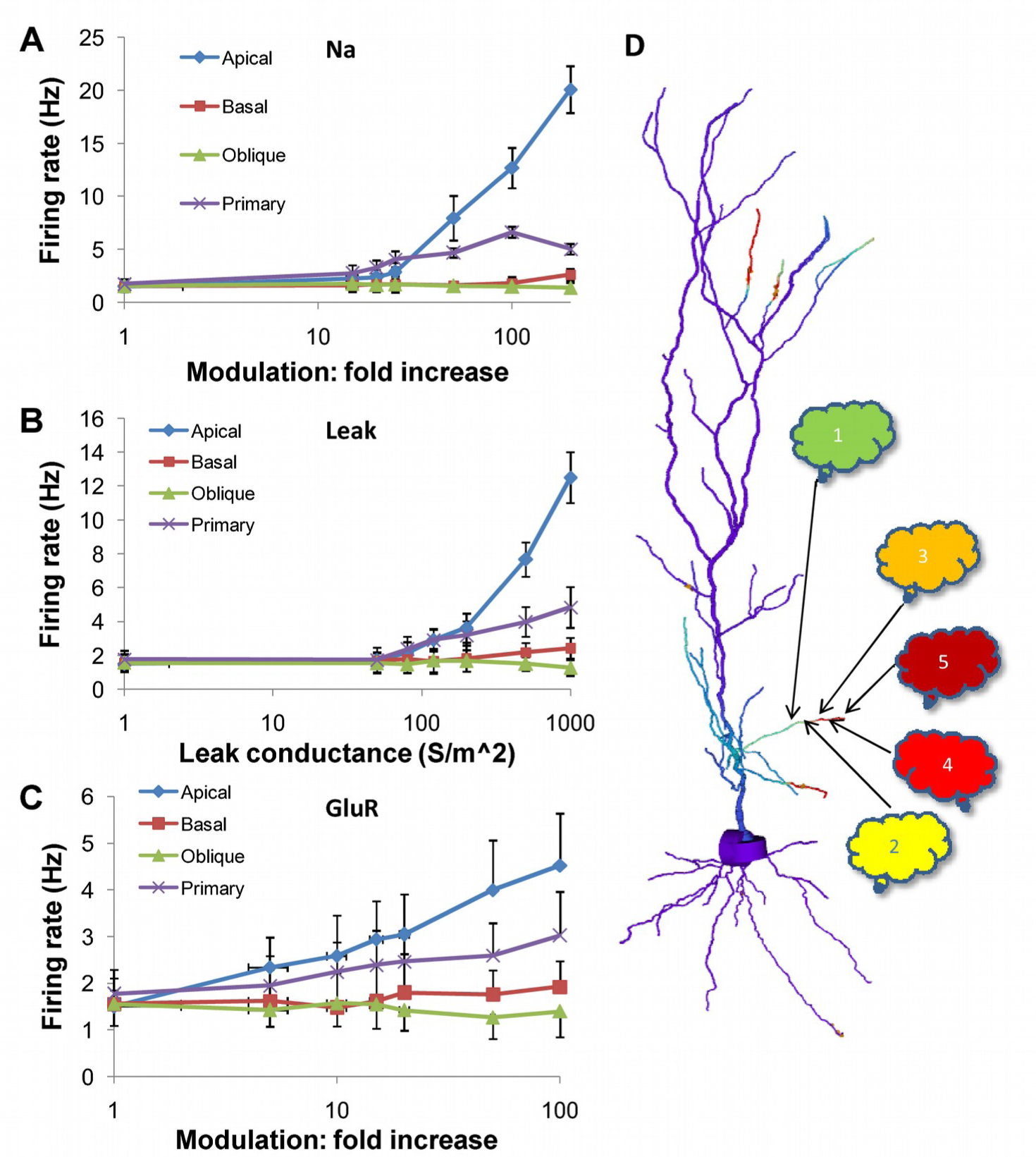
Firing rate changes in response to channel modulation in small zones on the dendritic tree. Error bars are standard deviation. A. Modulation of Na current. B. Adding Leak current. C. Modulation of GluR in the presence of the same synaptic input as was used as a stimulus for the sequential input. D. Schematic of convergence of inputs from different ensembles onto one dendritic zone. The numbered clouds represent input ensembles from which a single neuron projects to the indicated location on the dendrite.

We then asked if receptor modulation would be more effective if we factored in the strong synaptic input at the time of the last burst in the sequence. We repeated the calculations with this synaptic burst, and found that now a 10x increase in AMPAR conductance in the 20 micron zone in the apical dendrite led to a doubling in cell firing (Figure 7C). A similar manipulation in the primary dendrite had a shallower increase, about 1.5x.

Thus, chemical signaling, even in rather small regions of the apical dendrite, may lead to a rapid change in cell firing by local channel modulation. However, large channel modulations were needed to elicit sufficiently large (e.g., 50%) changes in in firing rate that could be detected over the noisy background.

## Discussion

We postulate that recursive sequence recognition is a fundamental computational operation in the brain, and that this operation may be implemented by neurons able to recognize spatially and temporally sequential input on behavioral time-scales (seconds). In this study we have examined three key parts of such a hypothesis. 1. At the chemical level, we have examined a set of abstract rate equations coupled by linear diffusion, and identified key motifs that support sequence recognition. 2. At the dendrite level, we have shown that these abstract principles of reaction-diffusion mediated sequence recognition carry over to much more detailed and biologically motivated neuronal models. These models are multiscale composite models, and explore the length-scales and physiological detail relevant to the current hypothesis. 3. At the cell level, we have shown that local, 20 micron channel modulation can elicit changes in somatic spiking. 20 microns is within the range of our predicted sequence-activated signaling.

### Sequence recognition confers substantial computational capabilities on single cells

Our model of sequence recognition in the neuron implements a potentially powerful form of cellular computation. It is interesting to estimate how much computation this represents. If we take a 5-stage sequence, a perfect sequence recognizer would distinguish 1 from 5! possible sequences, or 1 in 120. Longer sequences would have exponentially steeper discrimination ratios, but this places increasingly stringent constraints on network connectivity due to the requirement of having the inputs close to each other. Based on our simulations, biochemical reactions in the presence of noisy input are not as clean in their sequence selectivity as the abstract models (Figure 6 vs. Figure 3) but nevertheless do achieve good discrimination. In addition to being able to discriminate an ordered sequence from among many others, it is desirable that there be a large difference in chemical signal amplitude between ordered and scrambled sequences. Figures 2, 3 and 5 suggest that ratios in the range of 1:5 may be achievable, and 1:2 should be common.

Next, we estimate how many such sequence recognition blocks there might be in a neuron. For this analysis we draw on the results of the biochemical calculations for an estimate of a sequence recognition zone, of ~10 microns (Figure 6), and the result that sequence discrimination should happen over most of the dendritic tree except for the trunk of the primary apical dendrite (Figure 6J). Say the set of spines occupies 10 microns. The most conservative calculation requires that these blocks do not overlap, so in a pyramidal neuron with about 10000 microns of dendritic length we have ~1000 blocks. If we permit blocks to slide by one input at a time, there are 5 times as many sequence recognition blocks, i.e., ~5000. If the sequence lasts 10 seconds, a single block performs 0.1 sequence recognitions per second. Thus the neuron may perform 500 sequence discriminations each second. Calcium-induced Calcium release (CICR) based sequence logic (discussed below) is much faster, and may be able to increase this by a factor of 10.

A still more speculative calculation suggests that single-neuron sequence recognition may provide a way to handle the combinatorial explosion of possible input sequences. Assume that sequential inputs converge to 10 μm stretches of dendrite, having spine spacing *σ* = 0.5 μm. Assume that the sequence-recognition biochemistry tolerates a spatial slop of +-1.5 microns, or +-3 synapses, for each input. Then there are ~7 possible synaptic inputs for each stage of the sequence, and the overall block of 5 inputs may receive 7^5^, or ~17000 sequences of length 5. Using our calculation above, each neuron has ~5000 blocks, and so can recognize ~8.5×10^7^ sequences. Thus, with many assumptions about connectivity and sparsity, this mechanism of single-neuron sequence recognition suggests that single neurons may recognize very large numbers of input patterns through the combinatorics of synaptic convergence.

The key distinction between these two calculations is that the first indicates the computational capacity of a neuron, whereas the second considers the diversity of inputs it can act upon. Overall, our study suggests that sequence computation in the time domain, coupled with the combinatorics of spatially organized input along dendrites, result in extremely parallel and efficient computation at the single-neuron level.

### Single-cell sequence recognition may act on a continuum of timescales

Neural activity sequences occur at a range of timescales. A striking example of this is the mapping of behaviorally-driven place-cell sequences (seconds) to rapid, 100 ms time-scale replay sequences (Wilson and McNaughton, 1994). The distinctive aspect of our current analysis is that it works for slow behavioral time-scales of seconds, which is typically challenging for electrical network computations (but see (Barak and Tsodyks, 2006; Goudar and Buonomano, 2015)). Our analysis suggests that chemical mechanisms can also support faster sequence recognition on the same length-scales (Figure 4). One way this might be implemented in the cell could be a faster kinase cascade than MAPK, such as PKA or PKC (Bhalla, 2002). Another might be local CICR, which has already been proposed to support propagating wave activity in dendrites (Hagenston et al., 2008; Kapur et al., 2001; Larkum et al., 2003; Lee et al., 2016; Plotkin et al., 2013; Ross, 2012) These have faster dynamics, of the order of 100 microns/s. Further, dendritic Ca^2+^ imaging suggests that the length-scale of dendritic CICR is similar (~10-20 microns) to that envisaged for sequence recognition in our study (Hagenston et al., 2008; Kapur et al., 2001; Larkum et al., 2003). Simulation studies have previously suggested that CICR may be a mechanism for integration of inputs and resulting in graded, persistent activity (Loewenstein and Sompolinsky, 2003). This study has some parallels with our analysis, in particular the presence of a propagating wavefront of chemical (Ca^2+^) activation. However, in the earlier CICR model the wavefront propagation was modulated by the summed synaptic input to the neuron rather than local and specific sequential synaptic input, which is the basis of our study.

Still faster sequence recognition (~40ms) has been demonstrated in the electrical domain, where forward sequences can be discriminated from backward (Branco et al., 2010). This electrical mechanism has been shown to have ~40% discrimination between forward and backward patterns, rather than the strong selectivity among permutations of patterns shown by our chemical system. Thus this form of electrical sequence recognition is likely to operate in different network contexts than our proposed chemical recognition mechanism.

Together, we suggest that chemical and CICR-based sequence recognition may span the range of timescales from 200 ms to 10 s. It is interesting to speculate that these may coexist, thus permitting the same segment of dendrite to recognize slow behaviorally-driven sequences and also much faster replays of the same sequence.

### Plasticity is a likely target of biochemical sequence recognition

In the current study we have examined just one possible outcome of biochemical sequence recognition: the immediate impact on cell firing rate through channel modulation. However, several synaptically activated pathways may fit our analysis of state switching with feedback inhibition. These include four major kinases (PKA, PKC, MAPK, CaMKII) and a variety of second-messenger and metabotropic pathways (Bhalla and Iyengar, 1999; Kim et al., 2011; Lisman and Zhabotinsky, 2001). All of these have been proposed to play important roles in synaptic plasticity. We speculate that the large buildup of chemical activity during sequence recognition is well suited to play a role in various kinds of plasticity. It is intriguing to note that the current analysis of dendrite-based local chemical activation has parallels with the phenomenon of synaptic tagging (Frey and Morris, 1997). In both cases, synaptic input can amplify events in nearby, weakly-stimulated synapses. We speculate that sequence recognition and its conversion to plasticity events may share mechanisms with synaptic tagging.

### Sequence recognition is testable

Our hypothesis of single-cell sequence recognition is readily testable using current technology. Its chemical basis may be examined in brain-slice using local agonist application or sequential glutamate uncaging followed by microscopic readouts of activity reporters for candidate pathways. Such experiments have already been done to examine CICR triggered by synaptic inputs converging to adjacent synapses (Hagenston et al., 2008; Plotkin et al., 2013). These show that there is indeed a buildup of Ca^2+^ along small, 20 mm stretches of dendrite, under suitable stimulus conditions. If local biochemical signaling leads to changes in cellular firing-rate (e.g., Figure 7, (Plotkin et al., 2011)), then sequential uncaging along with patch recordings should report these. Further, pharmacological experiments in the slice would readily be able to tease apart possible mechanisms.

Another implication of the proposed reaction-diffusion mechanism for sequence recognition is that it suggests mechanisms for coupling genetic polymorphisms and mutations to a specific aspect of neuronal computation. For eample, small changes in reaction rates may shift the preferred timescale of recognized sequences (Figure 4), but large changes may eliminate tuning altogether. It would be interesting to see if there is a correlation between specific dendritic signaling genes and psychophysical measures of sequence computation.

In summary, we propose a novel computational function of single neurons, to recognize slow sequences on behavioral timescales, and to exploit combinatorics of input projections to carry out such recognition in a massively parallel manner. We have used simulations of abstract chemistry and detailed multiscale neuronal physiology to understand implications and constraints for such computation to occur. We suggest that this multiscale cellular signaling process may underlie computationally powerful sequence recognition mediated by single neurons.

## Methods

All modeling was carried out using MOOSE, the Multiscale Object-Oriented Simulation Environment (Ray and Bhalla, 2008). MOOSE is freely available and can be downloaded from moose.ncbs.res.in and GitHub. MOOSE utilizes the GNU Scientific Library (GSL) Runge-Kutta-Fehlberg 5^th^ order method for chemical computations, and a custom-written branching 1-D diffusion solver using the backward Euler (implicit) method. Stochastic chemical calculations were carried out using a custom-written optimized version of the Gillespie Stochastic Systems Algorithm. Pseudo-random numbers were generated by the Mersenne twister (Matsumoto and Nishimura, 1998). Electrical computations utilized a custom version of the branched nerve equation solution methods described by Hines (Hines, 1984). This has been validated by comparison with other simulators (Gleeson et al., 2010). Interfaces between chemical and electrical signaling components of the model utilized adaptor classes in MOOSE, which average over spatial and temporal discretization differences between the two methods. For example, electrical calculations (yielding Ca^2+^ values) utilize a much smaller timestep (~50 μs; 5 μm) but typically employ a larger spatial step than chemical calculations (~1 ms, 1 μm). The adaptors synchronized the chemical and electrical models every millisecond using first-order corrections to each model system. Electrical model morphologies were either geometrical (cylindrical) or derived from published cell reconstructions available on NeuroMorpho.org (Ascoli et al., 2007; Dougherty et al., 2012). Spines were positioned along dendrites using Poisson statistics with a specified mean spacing between spines (1 μm in the full cell model). Spines were modeled as a cylindrical head (0.5 μm length and diameter) on a cylindrical shaft (1 μm length, 0.2 μm diameter). In some simulations the spine dimensions were scaled up by 20% or 40%. Analysis and plotting was done using Python, NumPy, and MatPlotLib. Simulations were carried out on a variety of Linux workstations, and large calculations were carried out on Linux clusters. Figure generation code for Figures 2, 5 and 6 is available as supplementary material.

Chemical models are available in the supplementary material in tabular form and as GENESIS/kkit files, and will be hosted on http://doqcs.ncbs.res.in and ModelDB. Electrical models are available in supplementary material in tabular form and as MOOSE scripts. These scripts should be exact as of the GitHub version of 25 January 2017, but subsequent versions may have small output changes due to updates to the random number system. The morphology file is from NeuroMorpho.org (Ascoli et al., 2007; Dougherty et al., 2012) and attached in the supplementary material as the file VHC-neuron.CNG.swc.

## Author Contributions

USB is sole author.

## Acknowledgements

I acknowledge funding support from NCBS/TIFR and the Department of Science and Technology grant SR/CSI/66/2013 under the Cognitive Science Research Initiative. Large simulations were carried out in the NCBS Supercomputing facility. I acknowledge Arvind Kumar and Sahil Moza for comments on the project.

